# Production of Infectious Reporter Murine Norovirus by VP2 trans-Complementation

**DOI:** 10.1101/2023.07.27.550866

**Authors:** Ryoka Ishiyama, Kazuhiro Yoshida, Kazuki Oikawa, Reiko Takai-Todaka, Akiko Kato, Kumiko Kanamori, Akira Nakanishi, Kei Haga, Kazuhiko Katayama

**Author notes:** Address correspondence to Kei,HAGA, Laboratory of Viral Infection, Department of Infection Control and Immunology, Ōmura Satoshi Memorial Institute & Graduate School of Infection Control Sciences, Kitasato University, Tokyo, 108-8641; Japan, E-mail address, Kazuhiko KATAYAMA,Laboratory of Viral Infection, Department of Infection Control and Immunology, Ōmura Satoshi Memorial Institute & Graduate School of Infection Control Sciences, Kitasato University, Tokyo, 108-8641; Japan. These authors contributed equally to this work (Ryoka Ishiyama, Kazuhiro Yoshida, Kazuki Oikawa).

## Abstract

Human norovirus (HuNoV) causes gastroenteritis, a disease with no effective therapy or vaccine. Murine norovirus (MNV) easily replicates in cell culture and small animals and has often been used as a model to elucidate the structural and functional characteristics of HuNoV. A MNV plasmid-based reverse genetics system was developed to produce modified recombinant virus. In this study, we attempted to construct the recombinant virus by integrating a foreign gene into MNV ORF3 that encodes the minor structural protein VP2. We found that deletion of VP2 expression abolished infectious particles from MNV cDNA clones, and supplying exogenous VP2 to the cells rescued the infectivity of cDNA clones without VP2 expression. In addition, we found that the coding sequence of C-terminal ORF3 was essential for cDNA clones compensated with VP2 to produce infectious particles. Further, the recombinant virus with exogenous reporter genes in place of the dispensable ORF3 coding region was able to propagate when VP2 was constitutively supplied. Our findings indicate that foreign genes can be transduced into the norovirus ORF3 region when VP2 is supplied and that successive propagation of modified recombinant norovirus could lead to the development of norovirus-based vaccines or therapeutics.

**IMPORTANCE:** In this study, we revealed that some of the coding regions of ORF3 could be replaced by foreign gene and infectious virus could be produced under conditions with VP2 supplied. Propagation of this virus depended on VP2 being supplied *in trans*, indicating that this virus could infect only once. Our findings help to elucidate the functions of VP2 in virus lifecycle and to the development of other caliciviral vectors for recombinant attenuated live enteric virus vaccines or therapeutics.

## INTRODUCTION

Norovirus (NoV) is a positive-sense single-stranded RNA virus of the *Caliciviridae* family. Human norovirus (HuNoV) is the major causative agent of acute gastroenteritis, with more than one million cases worldwide each year (1). Although recent progress in *in vitro* HuNoV propagation with intestinal organoid-based systems provides hope for controlling and preventing HuNoV infections, no therapeutics or vaccines are yet available for this virus (2). However, much of the knowledge on noroviral infection was derived from studies using the murine norovirus (MNV). Its etiology is similar to that of HuNoV (3, 4), and most importantly, it can be efficiently propagated in tissue culture (5). Studies with MNV have provided a foundation for understanding the mechanistic details of the noroviral infection process (4, 6).

The noroviral genome encodes three open reading frames (ORFs). ORF1 encodes a nonstructural polyprotein that is self-cleaved by its own protease subunit to generate six proteins: Nterm (NS1/2), NTPase (NS3), 3A-like (NS4), VPg (NS5), protease (NS6), and RNA-dependent RNA polymerases (RdRp) (NS7). ORF2 and ORF3 encode the major capsid protein VP1 and the minor capsid protein VP2, respectively. The coding sequence of an additional ORF in the MNV genome, ORF4, overlaps with ORF2 encoding virulence factor 1 (VF1), but in different reading frames. These sequences are involved in MNV pathogenesis (7). Three regions in the MNV RNA genome form functional secondary structures that are important for viral propagation: the region overlapping with 5′ untranslated region (UTR) and ORF1 region, the region at proximal ORF2 start codon, and the region overlapping with the 3′ distal ORF3 region and 3′ UTR (8, 9).

The viral capsid, about 35 nm in diameter, is composed of 180 VP1 proteins that enclose the RNA genome and are linked at the 5′ end to VPg. The number of VP2 molecules in the virion is as-yet unclear (10–12). HuNoV VP2 is predicted to align with the inner surface of the VP1 shell and bind the VP1 dimer, which assumes that a virion can accommodate 90 VP2 proteins (11), whereas VP2 detection by western blotting of either the virion or VLP preparation revealed that few VP2 proteins were present in each particle (10, 12).

Noroviral VP2 is a highly basic RNA-binding protein (10, 11, 13). It is also highly phosphorylated upon overexpression in insect cells (13). HuNoV VP2 amino acid sequences are divergent (14), and their functional domains are poorly understood. Studies on VP2 in feline calicivirus, which belongs to the Vesivirus genus in *Caliciviridae*, are highly suggestive of the role of noroviral VP2 in viral infections. Structural studies using cryoelectron microscopy and tomographic analyses revealed that, upon binding of the capsid to the cellular receptor JAM1, the internal VP2s were aligned and extended outward to form portals (15). These might be for the exit of viral RNA and suggest that VP2 plays a critical role in the delivery of viral RNA in to host cells. A genetic approach to examine the VP2’s role in feline calicivirus using a viral mutant expressing VP2 showed that VP2 was essential in viral infection, and the lack of functional ORF3 could be complemented by co-expressing VP2 during the infection process (16).

The plasmid-based reverse genetics system enabled genetic engineering of the noroviral genome (17), but the genome regions that allow sequence modifications are not fully understood because of the presence of undiscovered functional sequence elements. Genetic modifications by random mutagenesis using transposon-mediated insertions revealed that the coding regions of NS4 and VP2 accept foreign sequence insertions (18). Thus, unlike other regions in the noroviral genome sequence, the VP2 coding region can accept insertion of short sequences, such as the FLAG tag; however, it is unclear if a foreign gene can be inserted into the VP2 coding region. In this study, we established a system to evaluate the RNA sequence region of ORF3, which is essential for the formation of the infectious particles, while complementing the VP2 protein *in trans*. Our findings allowed us to generate MNV reporter viruses carrying either fluorescent or luciferase genes.

## RESULTS

### Minor capsid protein, VP2, is required for forming infectious particles

Since ORF3 of MNV can accommodate the insertion of a FLAG tag (18), we examined VP2 function in virus infection to determine the essential portions of the VP2 region. To produce mutant MNV that does not express VP2 or that expresses a truncated-VP2, pMNV_S7F_ was applied. This plasmid was also used to produce progeny MNV for reverse genetic engineering (17). For constructing VP2-deleted virus, two stop codons were introduced at the 7^th^ and 10^th^ amino acid (pMNV_S7F_ORF3_stop_ (ORF3_stop_)) positions, and for several truncated VP2 expressing mutants, in-frame deletions of the VP2 coding sequence were constructed from 4^th^ to 105^th^ (pMNV_S7F_ORF3ΔN (ΔN)), from 53^rd^ to 156^th^ (pMNV_S7F_ORF3ΔM (ΔM)), and from 115^th^ to 205^th^ (pMNV_S7F_ORF3ΔC (ΔC)) (Fig. 1A). These plasmid constructs were transfected into 293T cells, and 48 hours later, the culture supernatant was transferred to RAW264.7 cells (Fig. 1B). The original construct, pMNV_S7F_, showed production of progeny infectious MNV particles since viral proteins (VPg and VP1) were expressed in RAW264.7 cells, whereas the constructs for VP2 protein deletion or each truncated-VP2 protein showed no viral proteins expression in RAW264.7 cells (Fig. 1C).

**Fig. 1.**
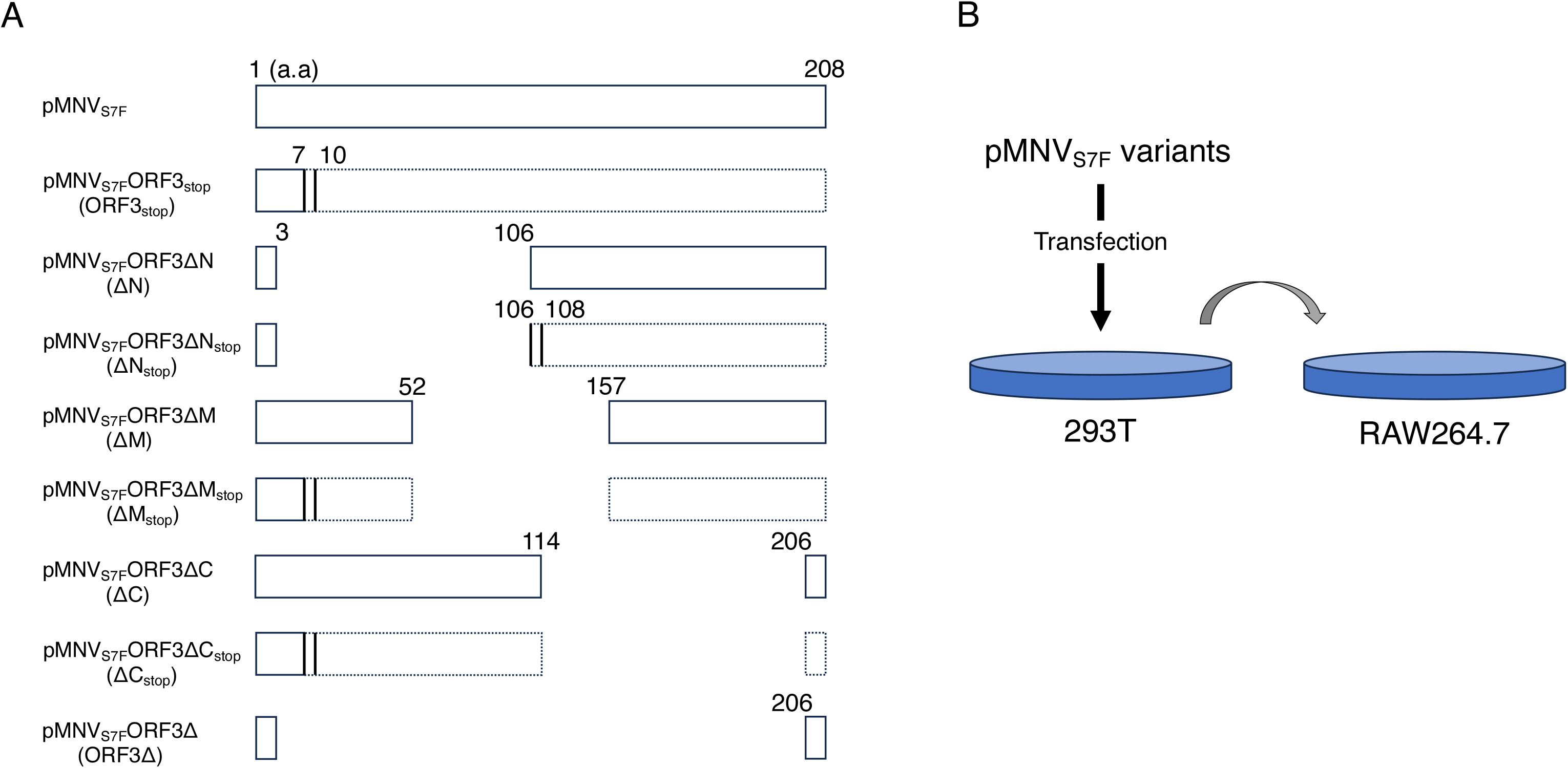
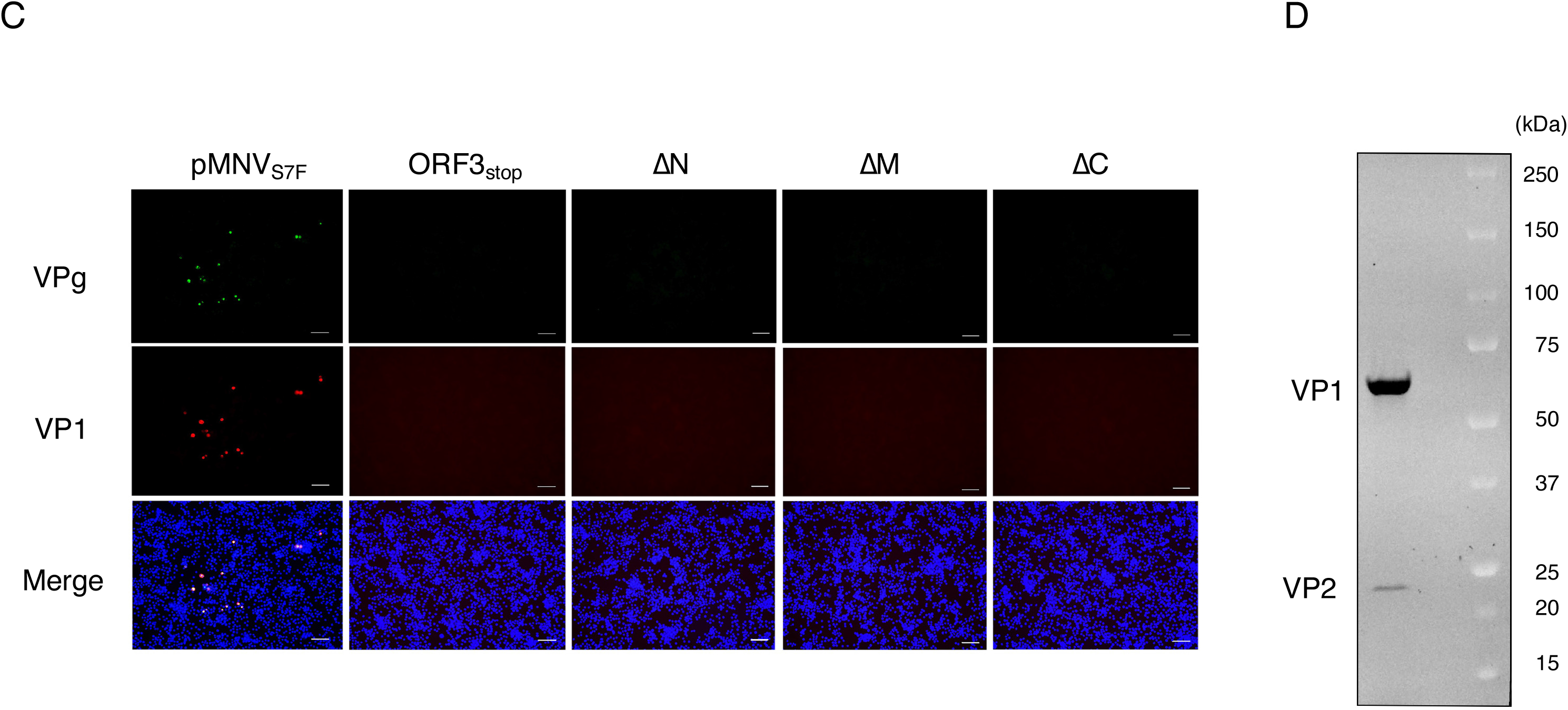
Deletion of VP2 protein prevented production of infectious MNV. A, Schematic diagram showing mutations and deletions in VP2 coding region of pMNV_S7F_ vector. Numbers indicate the position of the VP2 amino acid (a.a.) residues, and vertical lines with number indicate the position of the insertion of the in-frame stop codon. Coding region disrupted by stop codon was enclosed with dotted lines. B, Schematics of passages of MNV mutants. Each vector was transfected into 293T cells. After 48 hours of incubation, culture supernatant of transfected 293T cells was transferred to RAW264.7 cells. C, Examination of infectious MNVs produced by each construct. Detection of infectious events in RAW264.7 cells by the infection of viral mutants defective in expressing VP2. Each indicated construct was transfected into 293T cells, and after 48 hours incubation, each culture supernatant was transferred to RAW264.7 cells. After 24 hours of incubation, infectious events were detected by immunostaining for MNV VPg (green) and VP1 (red). The merged images were shown. Scale bars are 60 μm. D, Images obtained using the ChemiDoc MP imager. Purified and concentrated MNV was loaded onto Mini-PROTEAN TGX Stain-Free Protein Gels (Bio-Rad) and was electrophoresed. The band signal was obtained by detecting the fluorescence emitted by the reaction of the tryptophan residues of proteins with the excited trihalogenated compounds.

Although HuNoV and MNV virus-like particles were produced without VP2 (19–21), infectious MNV particles were required VP2 for infections and progeny production. Indeed, purified infectious MNV particles contained VP1 and VP2 (Fig. 1D); therefore, deletion or truncation of MNV VP2 disrupted the production of infectious particles. These findings indicated that deletions of approximately 100 amino acids in the ORF3 region prevented the formation of infectious particles, and so, it would be nearly impossible to insert foreign genes in VP2 coding region.

### Functional complementation by supplying VP2 in *trans*

Since VP2 was required to produce infectious particles, exogenous VP2 protein was expressed in 293T cells and RAW264.7 cells, and a continuous supply of VP2 was provided throughout the post-transfection of each construct. The culture supernatant of 293T co-transfected with pMNV_S7F_ORF3_stop_ and pORF3 for VP2 expression was transferred to RAW264.7 cells expressing VP2 (RAWVP2) (Fig. 2A and B). After 48 hours of incubation, VPg and VP1 were expressed in RAWVP2 cells (Fig. 2C, “pMNV_S7F_ORF3_stop_”), albeit a smaller number than in pMNV_S7F_ (Fig. 2C, “pMNV_S7F_”). This indicated a partial recovery of the infectious particles derived from ORF3_stop_ that lacked VP2 expression. In contrast, the construct lacking most of the ORF3 sequence (pMNV_S7F_ORF3Δ (ORF3Δ)) or lacking RdRp expression (pMNV_S7F_ΔRdRp (ΔRdRp)) could not be recovered by VP2 *trans* complementation (Fig. 2C).

**Fig. 2.**
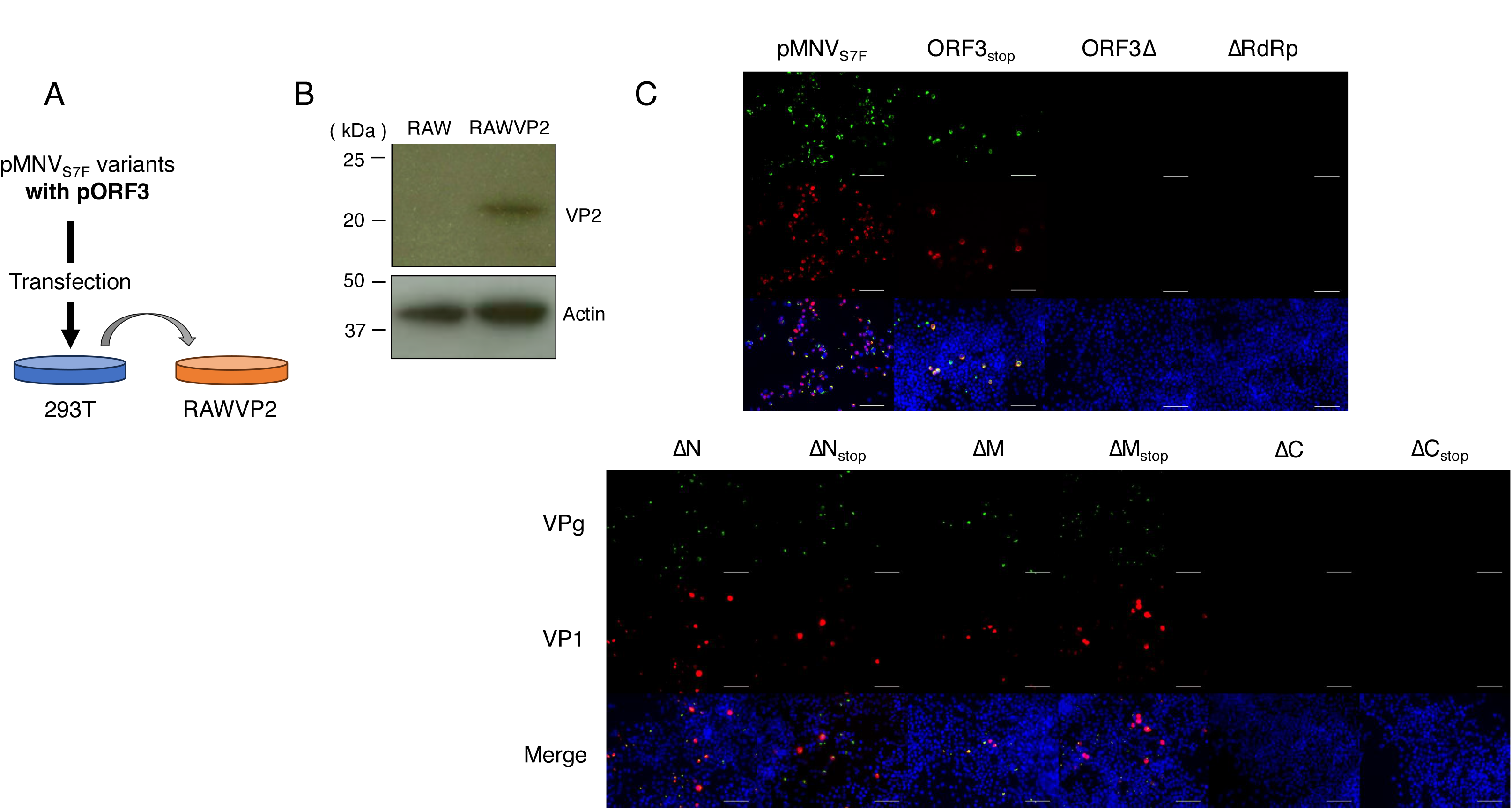
The sequence coding a.a. 114–205 of VP2 is essential for producing infectious MNV. A, Schematics of passages of MNV mutants. Each vector was co-transfected with pORF3 to express VP2 into 293T cells. After 48 hours of incubation, culture supernatants of transfected 293T cells were transferred to RAWVP2 cells. B, Western blotting to detect VP2 in RAWVP2. The positions of molecular mass were marked on the left. Actin visualized with an anti-antibody was used as a loading control. C, Examination of infectious MNVs produced by each construct. Detection of infectious events in RAWVP2 by the infection of viral mutants defective in expressing VP2. Each indicated construct was co-transfected with pORF3 in to 293T cells, and after 48 hours incubation, each culture supernatant was transferred to RAWVP2 cells. After 24 hours of incubation, infectious events were detected by immunostaining for MNV VPg (green) and VP1 (red). The merged images were shown. Scale bars are 60 μm.

Next, we determined the RNA coding sequences of ORF3 that were necessary for forming infectious particles under exogenous VP2 conditions. In addition to the ΔN, ΔM, and ΔC constructs, pMNV_S7F_ΔN_stop_ (ΔN_stop_), pMNV_S7F_ΔM_stop_ (ΔM_stop_), and pMNV_S7F_ΔC_stop_ (ΔC_stop_) were designed by introducing two stop codons immediately after each start codon to eliminate the truncated VP2 proteins (Fig. 1A). Infectious events were observed in RAWVP2 cells that were incubated with the products derived from co-transfection of ΔN, ΔM, ΔN_stop_ or ΔM_stop_ construct with pORF3 in 293T cells (Fig. 2C). Stop codon insertions to ΔN or ΔM did not significantly alter the efficiency of virus recovery, indicating that truncated VP2 protein from ΔN or ΔM did not impair the VP2 function supplied in *trans*. Meanwhile, exogenous VP2 expression could not rescue infectious particles from the ΔC and ORF3Δ constructs (Fig. 2C). The failure of infectious events from RAWVP2 cells was not due to expression of C-terminal lacking VP2 protein from the ΔC construct, because ΔC_stop_, which did not express truncated-VP2, also showed no infectious events in RAWVP2 cells. These findings indicated that residues 115–205 of VP2 are essential for producing infectious particles even with VP2 complementation.

### Genomic sequences encoding the VP2 C-terminal region are essential for infectious particle production

We also determined if infectious particles were produced from each construct upon VP2 complementation. Viral particles in the culture medium were immunoprecipitated with an anti-VP1 antibody, the RNA was purified, and the RNA copies of MNV genome were counted. If MNV genomic RNA was co-immunoprecipitated with VP1 protein, viral particles with genome RNA would be present in the culture supernatant. Copy numbers of viral RNA could be quantified by nested RT-PCR with a known copy number (Fig. 3A). Detection of co-immunoprecipitated viral RNA in culture supernatant of transfected-293T cells by anti-VP1 antibody was consistent with the results showing the infectious events in RAWVP2 cells (i.e., ORF3_stop_, ΔN, ΔN_stop_, ΔM, and ΔM_stop_) (Fig. 3B). Co-immunoprecipitated viral RNA was more abundant in RAWVP2 culture medium than in 293T cells, meaning more particles appeared to be produced after passaging (Fig. 3B, lanes 1 to 6). Clones ΔC (lane 7), ΔC_stop_ (lane 8), ΔORF3 (lane 9), and those defective in expressing RdRp (lane 10) yielded no detectable RNA by the immunoprecipitation with anti-VP1 antibody. In other words, they failed to form infectious particles in culture supernatant even in the presence of VP2 supply *in trans*. These findings suggested that the deficiency was due to a lack of particle formation rather than formation of non-infectious particles.

**Fig. 3.**
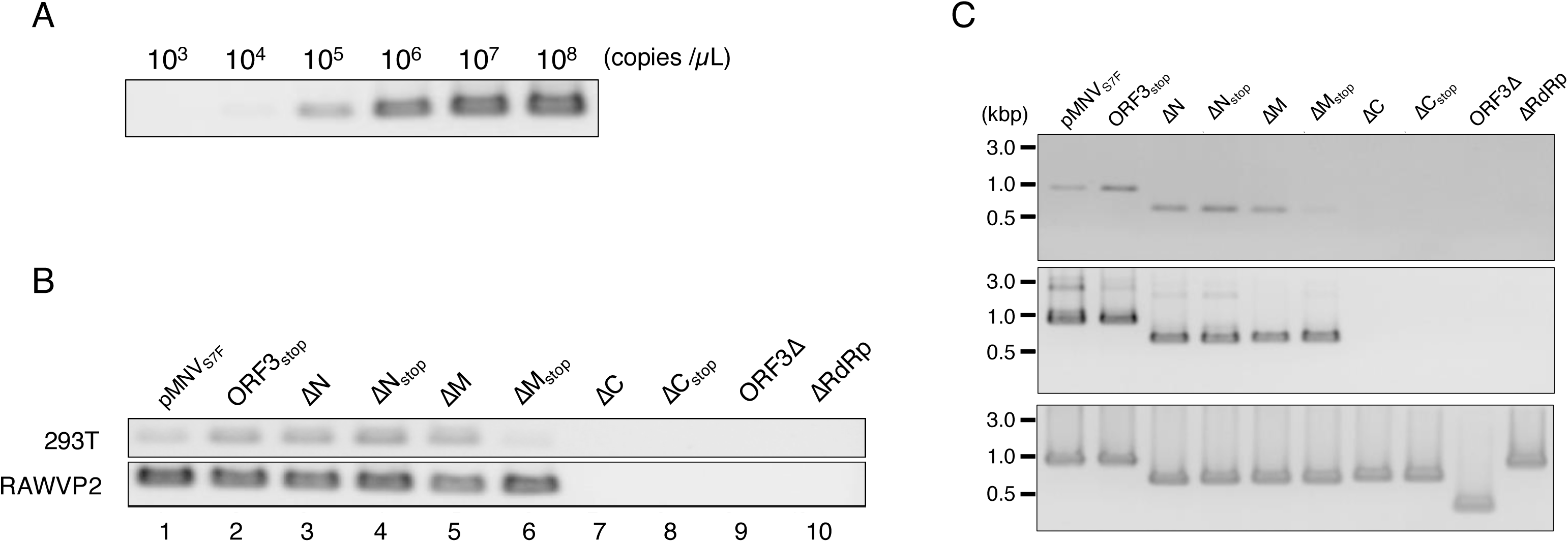
Detection of MNV RNA co-immunoprecipitated with anti-VP1 antibody from the cell-culture medium. A, Nested RT-PCR of standard RNA. Copies of MNV viral RNA synthesized *in vitro* were amplified by the nested RT-PCR as the reference to the signal detected. B, Detection of viral RNA co-immunoprecipitated using anti-VP1 in the cell-culture medium. Culture medium of 293T cells transfected with each viral construct with pORF3, and that of RAWVP2 was immunoprecipitated with anti-VP1 antibody and detected for viral RNA by nested RT-PCR. C, ORF3 deletions were maintained during propagation of the viral mutants harboring ORF3 mutations. The viral RNA immunoprecipitated with anti-VP1 antibody was amplified in ORF3 region by nested RT-PCR. Top panel; Amplified products from the culture medium of 293T cells co-transfected with each viral construct and pORF3 (“293T”). Middle panel; Amplified products from the culture medium of the RAWVP2 (“RAWVP2”). Bottom panel; PCR amplification of the ORF3 region from the viral molecular clones harboring mutations in ORF3 (“Plasmid”).

### Infectious events were not caused by revertants that reacquired functional VP2 coding region

The next step was to confirm that the observed infectious events were not due to revertants, which frequently occur (16). Thus, the length of the truncated-VP2 coding sequence was confirmed by amplifying the entire ORF3 region of each viral mutant by RT-PCR. Cell-culture medium, including the virus produced from the cells either transfected with the mutant viral constructs and pORF3 (Fig. 3C, “293T”) or infected in the presence of VP2 (Fig. 3C, “RAWVP2”), were immunoprecipitated with anti-VP1 antibody. Viral RNA was purified from the immunoprecipitates, and the ORF3 region was amplified. In agreement with the previous results, ORF3_stop_, ΔN, ΔN_stop_, ΔM and ΔM_stop_ appeared to generate viral particles carrying viral RNA, and passage in RAWVP2 yielded even more. By comparing these to the fragment size of the original plasmid construct (Fig. 3C, “Plasmid”), we found no obvious change in the ORF3 region of the mutants. This finding indicated that the introduced deletions were maintained during serial passage and no detectable revertants were generated during the experiments.

### Evaluation of VP2 complementation system using Huh7.5.1/CD300lf cells

To generate virus carrying a foreign gene, the MNV culture system was used for greater virus replication efficiency than RAW264.7. Huh7.5.1 cells, a subline of Huh7 hepatocarcinoma cell line, have a missense mutation in the RIG-I gene and are highly permissive to JFH-1 strain of hepatitis C virus (22). A MNV receptor molecule, mouse CD300lf, was transduced into Huh7.5.1, and highly efficient clonal cells were obtained after single-cell cloning. In agreement with published reports that used similar cells (23), Huh7.5.1/CD300lf cells were fully permissive to MNV and could produce higher amount of MNV than RAW264.7 cells (Fig. 4A). Thus, we expected more viruses to be produced by our modified reverse genetics system in this study.

**Fig. 4.**
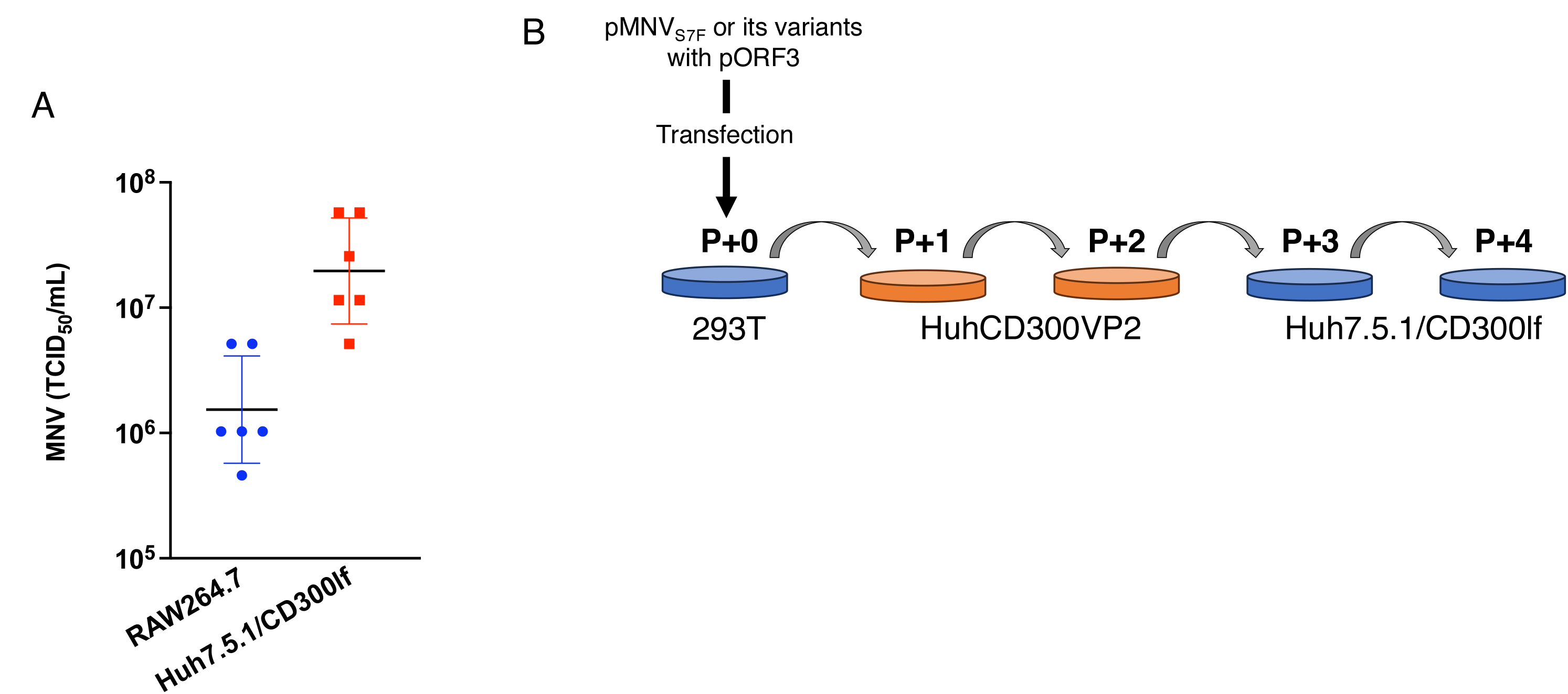
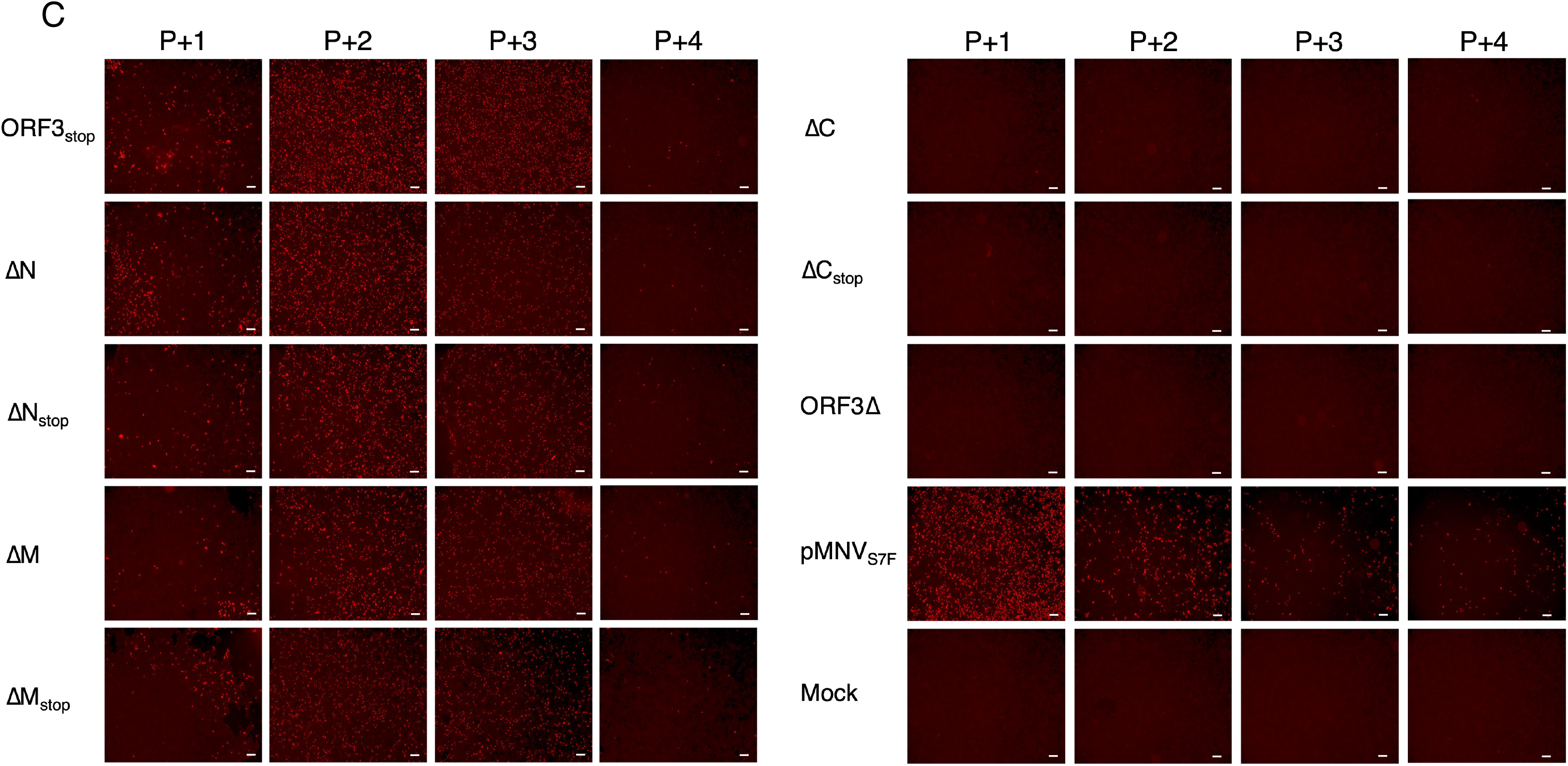
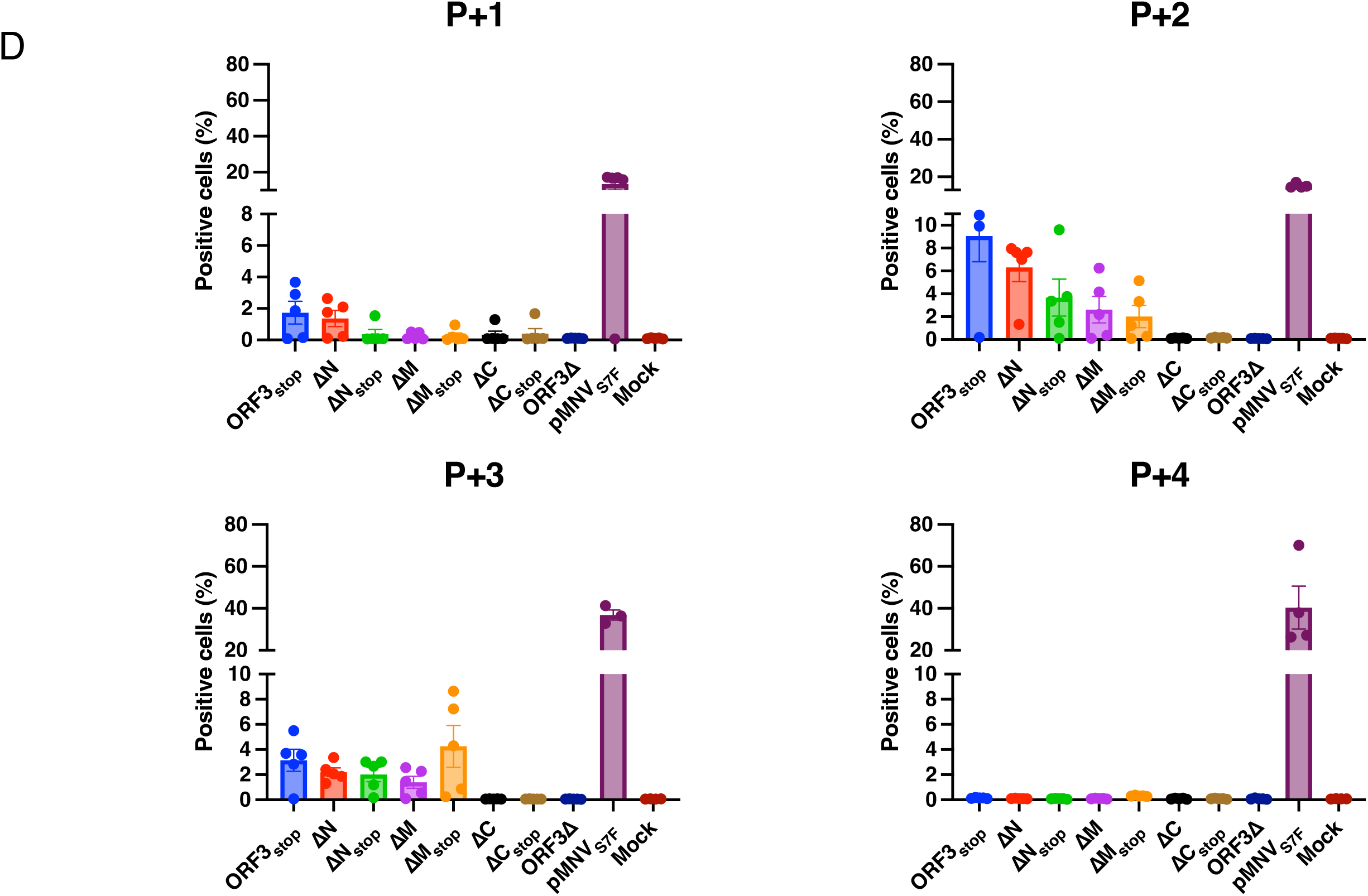
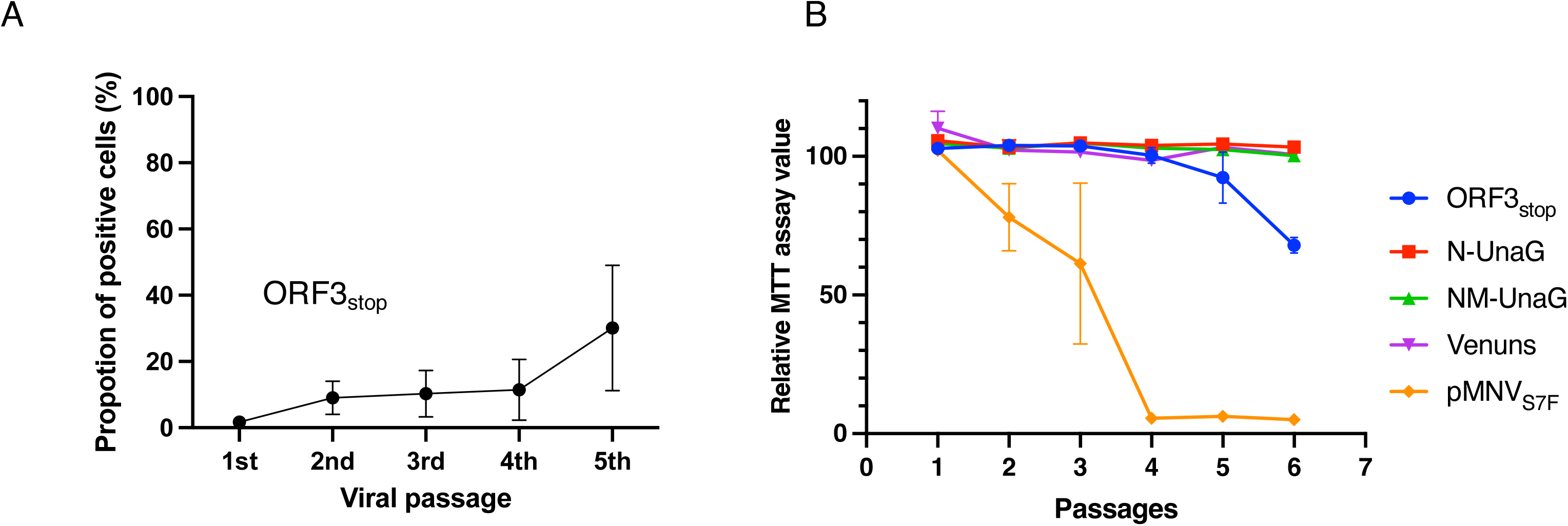
Huh7.5.1-based VP2 complementation system for propagating MNV VP2-defective mutants. A, The infectivity of wild-type MNV produced from RAW264.7 or Huh7.5.1/CD300lf cells was determined by 50% tissue culture infectious dose (TCID_50_) assay. Each cell line was infected with MNV (moi 0.1) and was incubated for 24 hours. The infectivity of the virus in each supernatant was determined by TCID_50_ using RAW264.7 cells. B, Schematics of passages of MNV mutants. Either MNV wild-type or mutants defective in expressing VP2 generated by transfection to 293T cells (293T) were infected into HuhCD300VP2 cells (P+1). The supernatant of P+1 was passaged in HuhCD300VP2 cells (P+2) and then twice in Huh7.5.1/CD300lf (P+3 and P+4). C, Representative images of HuhCD300VP2 (P+1 and P+2) or Huh7.5.1/CD300lf (P+3 and P+4) infected product from cell-culture medium of 293T cells transfected with pORF3 plus each construct. Cells in each passage was detected for infectious events by immunostaining of MNV N-terminal protein (red). For each construct and passage, whole image of each well was taken, and N-terminal protein positive cells were counted in five independent wells. Scale bar (200μm) in white are indicated. D, N-terminal protein-positive HuhCD300VP2 cells (P+1 and P+2) or Huh7.5.1/CD300lf cells (P+3 and P+4) were counted from whole image of each well using the built-in image processing software of the BZ-X 800. The percentage of infected cells was calculated by dividing the number of cells expressing the N-terminal protein by the total number of DAPI-positive cells. Each dot represents the percentage of positive cells in each image and the bar indicates the median value. All images used for measurements were shown in Supplementary Fig. 1.

Culture supernatants of 293T co-transfected with each construct with pORF3 for VP2 expression were transferred to Huh7.5.1/CD300lf expressing VP2 (HuhCD300VP2) (Fig. 4B), and cells expressing the N-terminal protein were counted. At the first transfer to HuhCD300VP2 cells (P+1), N-terminal protein expression was observed from ORF3_stop_, albeit a smaller percentage than in pMNV_S7F_ (Fig. 4C and D (P+1)). After another passage in the presence of VP2, the number of positive cells increased (Fig. 4C and D (P+2)). These findings indicated that partial recovery of the infectious particles derived from ORF3_stop_ was also observed in HuhCD300VP2, and the virus was propagated. We also counted the cells expressing viral proteins to determine if other mutant constructs produced infectious particles when VP2 was supplied. Infectious events were observed at the first transfer to HuhCD300VP2 cells that were incubated with the products derived from co-transfection of ΔN, ΔM, ΔN_stop_, or ΔM_stop_ construct with pORF3 in 293T cells (Fig. 4C and D (P+1)). Further passages to other HuhCD300VP2 cells increased the number of cells positive for viral protein (Fig. 4C and D (P+2)). However, the number of positive cells was lower when the cells were passaged to HuhCD300lf that was not supplied with VP2, and another passage yielded no positive cells (Fig. 4C and D (P+3 and P+4)), indicating that infectivity of this mutant virus depended on VP2 complementation. Meanwhile, VP2 complementation could not recover infectious particles provided from ΔC and ΔC_stop_, which was same result observed in RAWVP2 cells (Fig. 4C and D), and wild-type MNV (WT) derived from pMNV_S7F_ could keep their infectivity without VP2 complementation. The higher percentage of infected cells with WT was due to a reduction of total cell number by significant cell death.

### Production of reporter virus using VP2 complementation system with Huh7.5.1/CD300lf cells

Since passaging the virus from ORF3_stop_ in HuhCD300VP2 cells increased the number of cells expressing viral proteins, we repeated the passage of the virus among HuhCD300VP2 cells every 48 hours. After the 4th passage, about 40% of the cells were positive (Fig. 5A). As the number of positive cells increased, the viability of the infected HuhCD300VP2 cells decreased; however, the loss of viability was much less than in the WT virus derived of original pMNV_S7F_ vector (Fig. 5B), indicating that mutant virus production with exogenous VP2 was less inefficient than WT.

**Fig. 5.**
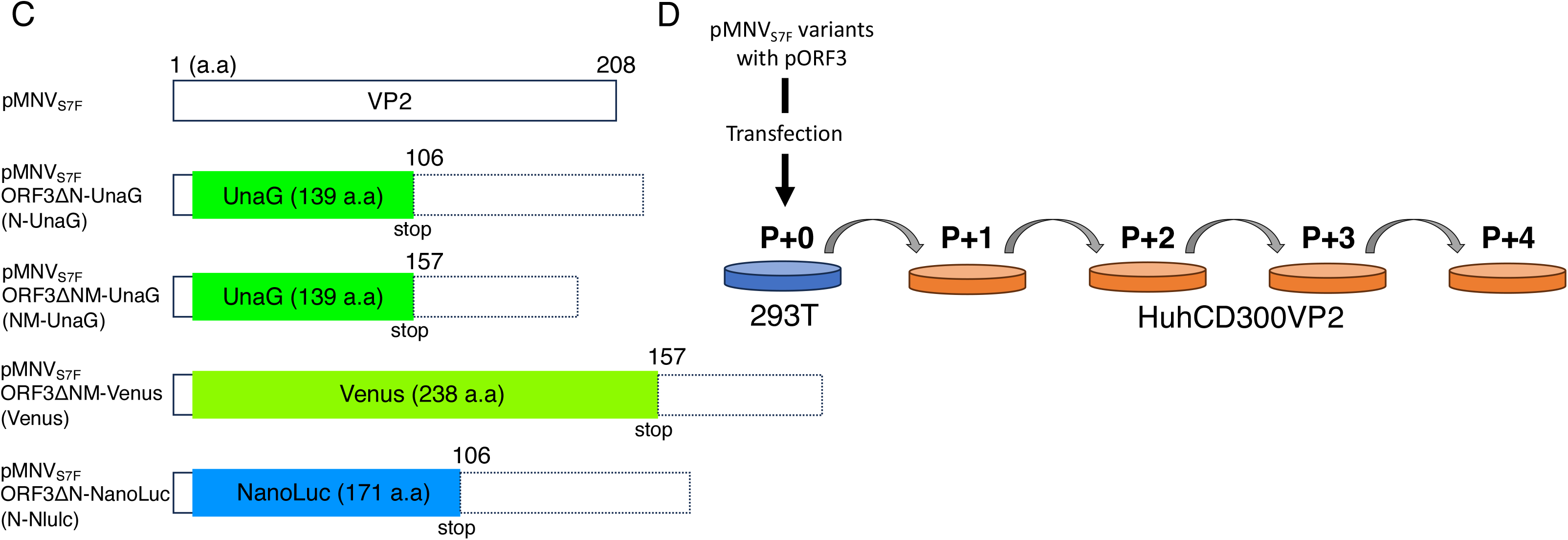
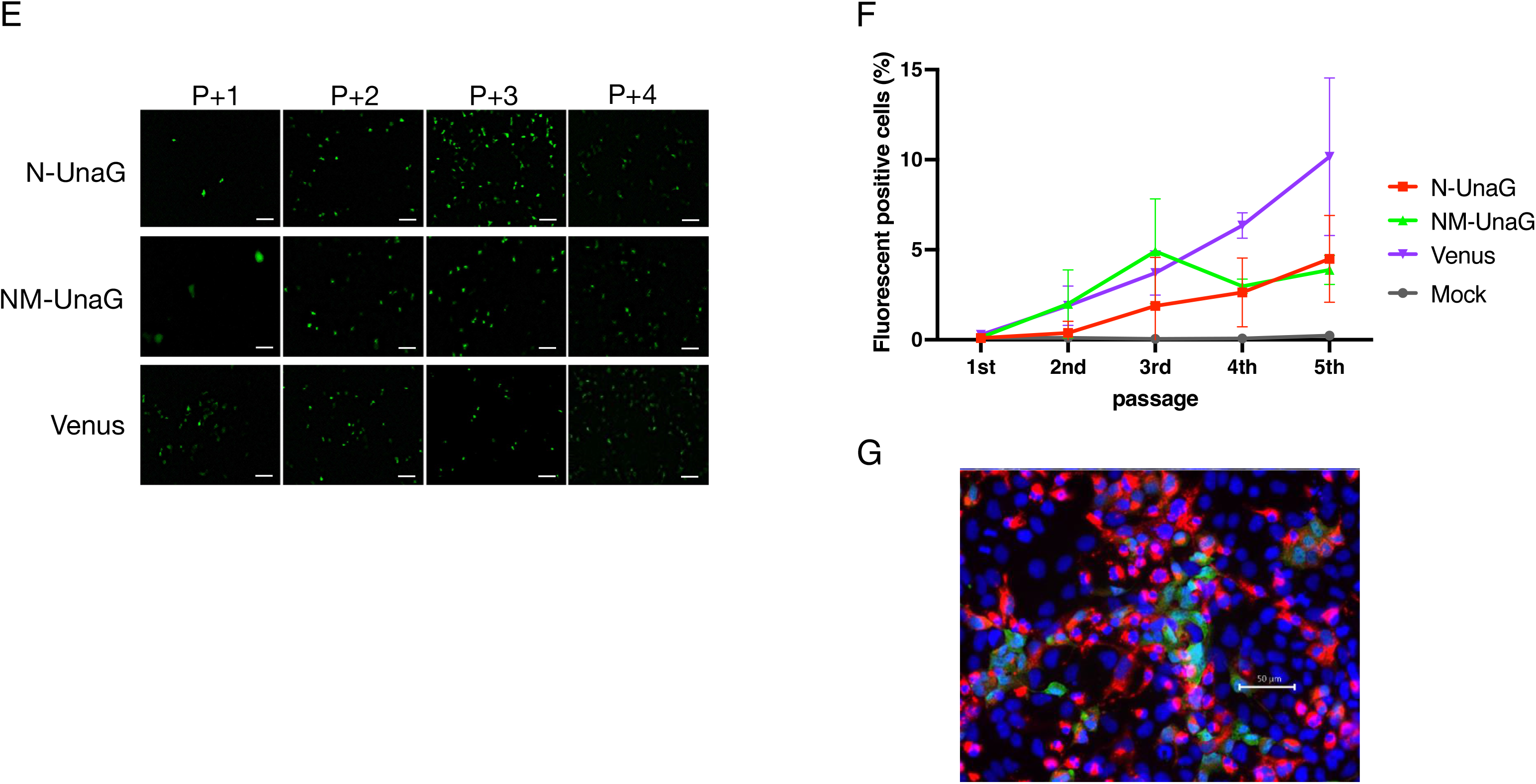
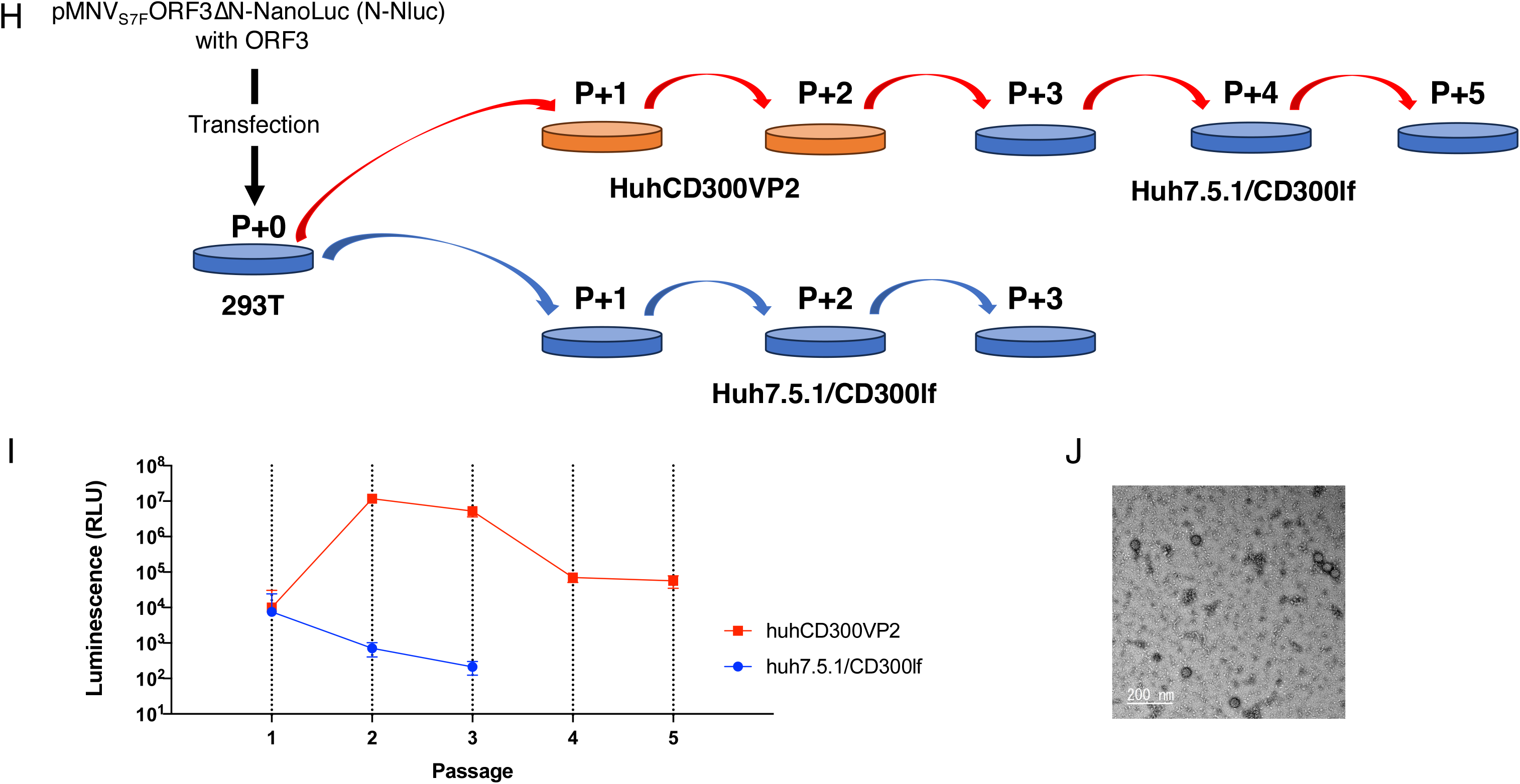
Propagation of MNV reporter virus in HuhCD300VP2 cells. A, The mutant generated by co-transfection of pORF3 and ORF3_stop_ was passaged in HuhCD300VP2 cells. B, Cell viability upon serial passages of MNV wild-type, VP2-defective mutant, and reporter viruses. HuhCD300VP2 was incubated with supernatant collected from each passage for 48 hours, and the extent of the formazan product converted from MTT by the cellular enzyme was examined by measuring OD_490_ value. “Relative MTT assay value” indicates the proportion of the sample MTT assay value divided by that of the control cells that did not receive viruses at each time-point. C, Schematic diagram of the reporter gene insertion into the VP2 coding region. Constructs used to generate the reporter viruses are listed at the left. Coding regions of reporter genes, UnaG and Venus, are highlighted in green, and NanoLuc was highlighted in blue. Numbers on the column indicate the position of VP2 amino acids. Boxes with dotted line indicate untranslated VP2 coding region due to the stop codon of the reporter gene. D, Schematic of the passages of MNV reporter mutants. MNV reporter mutants generated by transfection to VP2-expressing 293T cells (293T) were infected into HuhCD300VP2 cells (P+1). The supernatant of P+1 was repeatedly passaged in HuhCD300VP2 cells (P+2 to P+4). E, Serial passages of the reporter viruses in HuhCD300VP2 cells. Reporter constructs (“N-UnaG”, NM-UnaG” and “NM-Venus”) were co-transfected with pORF3 to 293T cells, and the culture media the transfected cells were used as the source of virus. Vertical axis indicates the proportion of cells positive for viral N-terminal protein detected by immunofluorescent technique (“ N-UnaG”, NM-UnaG” and “NM-Venus”). Scale bar (150μm) in white are indicated. F, The reporter viruses, N-UnaG, NM-UnaG, and NM-Venus, were serially passaged in HuhCD300VP2 cells, and cells positive for the fluorescent signal were detected with an epifluorescent microscope. Cells with the positive signal were counted, and the proportions of the positive cells are shown in Fig. 5E. G, Cells were infected with the NM-Venus reporter virus, fixed with 4% paraformaldehyde, and detected for viral N-terminal protein (red) and Venus fluorescence (green). Nuclei were stained with DAPI (blue). H, Schematics showing the method of production and passages of MNV reporter virus expressing NanoLuc luciferase (N-Nluc). Briefly, N-Nluc virus was produced by transfection to 293T cells with N-NLuc and pORF3. The supernatant of the transfected 293T cells (P+0) was applied to HuhCD300VP2 cells (P+1, red), and the supernatant was further passaged to HuhCD300VP2 (P+2, red). The supernatant was then applied to Huh7.5.1/CD300lf cells (P+3, red), and its supernatant was subsequently applied to Huh7.5.1/CD300lf (P+4 and P+5, red). The supernatant of the transfected 293T cells was also used to infect Huh7.5.1/CD300lf (P+1, blue). The supernatant of Huh7.5.1/CD300lf was used to passage Huh7.5.1/CD300lf (P+2 and P+3, blue). HuhCD300VP2 and Huh7.5.1/CD300lf were incubated with the supernatant collected at each passage for 72 hours. I, NanoLuc activity was detected at each passage upon infection with N-Nluc reporter virus. At each “Passage” depicted in H, cells were taken and assayed for NanoLuc luciferase activity using Nano-Glo® Luciferase Assay System (Promega). The vertical axis of the graph shows luminescence signal detected by luminometer, and the horizontal axis shows the number of passages. The red and blue lines depict data of passages in red and in blue (see H), respectively. J, Electron micrograph of N-Nluc reporter virus particles. Scale bar (200 nm) in white is indicated.

Our results consistently showed that a part of the VP2 coding region from the 4^th^ to 105^th^ or 53^th^ to 156^th^ residues was not needed if VP2 was supplied *in trans*. We then replaced that region with a fluorescent protein-coding sequence and determined if a virus with a modified viral genome could propagate in VP2-expressing cells and function as a reporter virus. We constructed three constructs carrying either UnaG or Venus genes (Fig. 5C). UnaG is one of the smallest fluorescent proteins whose fluorescence depends on the binding of bilirubin to fluorochromes (24). Venus is a variant of GFP that has been modified for better fluorescent properties (25). N-UnaG or NM-UnaG had the UnaG gene inserted between the 4^th^ and 105^th^ or 156^th^ residues, respectively, and NM-Venus had a Venus gene inserted between the 4^th^ and 156^th^ residues of the VP2 coding region (Fig. 5C). Each construct was co-transfected with pORF3 into 293T cells to generate the reporter virus and, subsequently, passaged in HuhCD300VP2 cells (Fig. 5D). Upon reporter virus infection, we observed fluorescent signals in some HuhCD300VP2 cells that likely expressed UnaG or Venus (Fig. 5E). Upon passages of ORF3_stop_ viruses in HuhCD300VP2 cells, nearly 40% of cells were positive for the viral antigen upon 5^th^ passage of VP2-defective mutants (Fig. 5A). In contrast, after passages of the reporter viruses, fluorescence was detected in fewer than 10% of cells (Fig. 5F). Simultaneous detection of viral antigen (N-terminal protein) and Venus protein revealed that the cells visibly expressing Venus protein were underrepresented among the cells expressing viral antigen (Fig. 5G).

Our reporter virus system accommodated a relatively large Venus gene, 714 bp in length, which led us to test the effectiveness of a 513-bp gene encoding a small luminescent protein, NanoLuc. The Nluc reporter virus was constructed by inserting the NanoLuc coding sequence in-frame between the 4^th^ and 105^th^ amino acid-coding regions of VP2 (N-Nluc) (Fig. 5C). The N-Nluc construct was co-transfected with pORF3 into 293T cells, and the resulting particles were used to infect either HuhCD300VP2 or Huh7.5.1/CD300lf cells (Fig. 5H). Upon N Nluc viral infection, a luminescent signal was evident in HuhCD300VP2 cells, but only a marginal signal was detected in Huh7.5.1/CD300lf cells (Fig. 5I). Propagated virus in HuhCD300VP2 cells disappeared after repeated passages in Huh7.5.1/CD300lf cells since the luminescent signal was abolished when the virus was passaged twice in Huh7.5.1/CD300lf. These findings indicated that Nluc expression depends on the presence of functional VP2. The reporter virus propagated in HuhCD300VP2 and particles resembling MNV virions were readily detected in the electron micrograph (Fig. 5J and Supplementary Fig. 2). Our results indicate that MNV-based reporter viruses carrying reporter genes inserted into VP2 coding region were able to propagate in cells when VP2 was supplied *in trans*.

## DISCUSSION

We used a genetic approach to examine the role of VP2 protein in MNV infection. The needs for VP2 protein and for the RNA sequence in ORF3 for a productive MNV infection were analyzed separately. First, a molecular clone of viral mutants was generated that lacked VP2 expression due to insertion of a stop codon immediately after first methionine of ORF3 (17). No infections were noted after transferring culture supernatant of 293T cells transfected with the RGS construct not expressing VP2 protein. However, supplying VP2 protein exogenously in 293T cells and subsequent MNV permissive cells restored infectious viral production. Second, several molecular clones with a deletion of the ORF3 RNA sequence were transduced into 293T cells expressing VP2, and we determined that the coding sequence of C-terminal (a.a. 115–205) of VP2 protein was essential for infectious virus production when VP2 was supplied in *trans*. Viral RNA immunoprecipitated with anti-VP1 antibody in the infected cell culture medium confirmed that the infectious events were mediated by viral particles. Moreover, the apparent propagation of the mutant was not due to revertants: no such events were observed when VP2 was not supplied *in trans* (Fig. 4B, C and D), and the nucleotide alterations were maintained during the passage of the mutant viruses (Fig. 3C). With HuhCD300VP2 cells, viral variants with genes encoding fluorescent or luminescent proteins were propagated in VP2-expressing cells and transduced to express the reporter genes in target cells. Thus, our study revealed some roles of VP2 in the viral infection cycle by exploiting the VP2 coding region and showed its potential as a noroviral reporter vector.

Approximately two-thirds of the proximal region of the VP2 coding region was not essential for viral propagation as long as VP2 was supplied *in trans*. Deletion of the C-terminal region of ORF3 was detrimental to viral lifecycle, even in the presence of VP2. These results are consistent with previous findings in feline calicivirus (FCV), where VP2 supplied *in trans* could not rescue deletion of the C-terminal VP2 coding region (16). The region encoding the C-terminal VP2 of MNV overlaps with a region predicted to form three stem-loop RNA secondary structures (8). The first stem loop, SL1, overlaps with the coding region of VP2 amino acid 166 to the end and was deleted in ΔC. Supplying VP2 *in trans* did not rescue the infection process of ΔC or ΔC_stop_, providing experimental evidence that the region encompassing the SL1-forming sequence is critical for the viral lifecycle. This had not been determined in previous studies (8, 9, 26).

A structural study of FCV entry suggested a role for MNV VP2 in the infection process. Upon FCV recognition by the cellular receptor JAM1, VP1 undergoes a conformational shift to expose the internal VP2, which self-assembles and forms a portal to release viral RNA from the viral capsid into the host cell (15). Note that the VP2s of FCV and MNV differ considerably in size and sequence. It is yet to be determined if the VP2 of MNV has a role in viral entry similar to that of FCV. Alternatively, the lack of VP2 could compromise the structural integrity of the viral particles because VP2 contributes to the stability of the capsid structure (27).

Assembly of the capsid is governed by VP1 and is carried out without VP2 and the viral RNA genome (19, 20). However, VP1 and VP2 directly interact in Norwalk virus (28) and FCV (15, 29), and in this study, purified MNV retained VP2 protein within virus particles (Fig. 1D). When VP2 was deleted in FCV, genomic RNA replication was not impaired in CRFK cells, but infectious progeny was not produced (16), indicating that FCV VP2 was involved in successful capsid assembly. Therefore, VP2 of MNV also may be involved in efficient assembly or RNA incorporation into viral particle.

Huh7.5.1-derivative cells carrying MNV VP2 performed better, and approximately 40% of cells were infected upon serial passaging (Fig. 5A) and with evident cytopathic effects (Fig. 5B). With the success of VP2 complementation in the Huh7.5.1-based system, the mutant reporter virus was propagated in HuhCD300VP2 cells. Each reporter gene was replaced with the 5′ part of the VP2 coding sequence, and these viruses grew with VP2 complementation in the Huh7.5.1-based system. The expression of fluorescent and luminescent proteins in this system suggests that it has potential as a reporter virus system. This system could be used to study various aspects of MNV and could also benefit the approach for generating HuNoV reporter vectors, which is of great importance for attaining valuable insights into the biology of HuNoV, the etiological agent of the most frequent cause of viral gastroenteritis. Further, if our findings apply to other calicivirus, like human norovirus or sapovirus which can infect to human, it could lead to develop an attenuated live viral vaccine that can be amplified using VP2 expressing cell lines and be expected to infect only once *in vivo* where VP2 is not supplied.

## MATERIALS AND METHODS

### Cell lines

The cell line, 293GP, from RIKEN CELL BANK, was maintained in Dulbecco’s Modified Eagle Medium (DMEM; Wako), supplemented with 10% fetal bovine serum (FBS) and MEM non-essential amino acid solution (Thermo Fisher Scientific). 293T, RAW264.7, and Huh7.5.1 (a gift from Dr. Francis V. Chisari) cells were maintained in DMEM supplemented with 10% FBS.

### Construction of plasmids

MNV expression vectors were constructed from pMNV_S7F_ (17) that harbored the MNV S7 strain cDNA (NCBI accession number AB435515). The pORF3 was constructed from pMNV_S7F_ by deleting ORF1 and most of the ORF2 coding regions.

VP2 mutations or deletions were introduced into the MNV cDNA of pMNV_S7F_. The 7^th^ (GGA) and 10^th^ (GGA) codons of ORF3 were changed to stop codons (TGA) to generate pMNV_S7F_ORF3_stop_ (ORF3_stop_). pMNV_S7F_ORF3ΔN (ΔN), pMNV_S7F_ORF3ΔM (ΔM), pMNV_S7F_ORF3ΔC (ΔC), and pMNV_S7F_ORF3Δ (ORF3Δ) were constructed by deleting ORF3 regions encoding 4–105, 53–156, 115–206, and 4–207 amino acids (aa), respectively. The constructs, pMNV_S7F_ORF3ΔN_stop_ (ΔN_stop_), pMNV_S7F_ORF3ΔM_stop_ (ΔM_stop_) and pMNV_S7F_ORF3ΔC_stop_ (ΔC_stop_), were made from ΔN, ΔM, and ΔC, respectively, by inserting two stop codons (TGA) in place of the 4^th^ (106^th^) and 6^th^ (108^th^) codons for pMNV_S7F_ORF3ΔN_stop_ and 7^th^ and 10^th^ codons for the remaining two constructs.

The pMNV_S7F_ORF3ΔN-UnaG (N-UnaG) and pMNV_S7F_ORF3ΔN-NanoLuc (N-Nluc) were made by replacing the ORF3 region encoding 5–105 amino acids with an in-frame insertion downstream of 4^th^ VP2 codon sequence of UnaG and NanoLuc (Promega) coding regions with their own stop codons, respectively. Similarly, pMNV_S7F_ORF3ΔNM-UnaG (NM-UnaG) and pMNV_S7F_ORF3ΔNM-Venus (NM-Venus) were made by replacing the ORF3 region encoding amino acids 5–156 with an in-frame insertion downstream of 4^th^ VP2 codon sequence of UnaG and Venus coding regions with their own stop codons, respectively.

MSCV-VP2-IRES-GFP, a mouse stem cell virus (MSCV) vector expressing VP2, was constructed from MSCV IRES GFP vector (Addgene #20672) by inserting the MNV VP2 coding region proximal to the internal ribosome entry site (IRES). pLVSIN-MNV VP2-IRES-puro was constructed to create a lentiviral vector expressing VP2 from pLNSIN-IRES-puromycin (30) by inserting the MNV ORF3 region.

The sequences of all the DNA constructs were confirmed by dideoxynucleotide sequencing (SeqStudio Genetic Analyzer, ABI) and next-generation sequencing (iSeq, Illumina).

### Production of RAWVP2, Huh7.5.1/CD300lf, and HuhCD300VP2 cells

RAWVP2 cells were generated by transducing the MSCV vector expressing MNV VP2 into RAW264.7 cells. The MSCV vector was produced by transfecting MSCV-VP2 IRESeGFP and VSV-G into 293GP cells using the Lipofectamine 3000 reagent (Thermo Fisher Scientific). At 48 hours after the transfection, the supernatant of 293GP containing the retroviral vector was collected and added to RAW264.7 cells, followed by incubation for 48 hours to generate RAW264.7VP2, herein RAWVP2.

Huh7.5.1/CD300lf cells were obtained from the Huh7.5.1 cells. Briefly, a lentiviral vector expressing CD300lf was made by co-transfecting pLVSIN-mCD300lf-IRES-hyg, which is a derivative of pLVSIN-mCD300lf-IRES-puro (30), with a hygromycin-resistance gene replacing that of puromycin, and three plasmid constructs, pMDLg/pRRE HIV-1, pMD2G env, and pRSV-Rev (Addgene), into 293T cells. This lentiviral vector was used to transduce the CD300lf expression cassette into Huh7.5.1 cells. The resultant Huh7.5.1/CD300lf cells were selected with 0.2 mg/mL hygromycin, and each single clone isolated by cell sorter (FACSMelody, BD) was tested for its ability to support MNV propagation. Meanwhile, a lentiviral vector expressing MNV VP2 was created by transfecting the constructs, pMDLg/pRRE HIV-1, pMD2G env, pRSV-Rev, and pLVSIN-MNV VP2-IRES-puro into 293T cells. Huh7.5.1/CD300lf clone12-2, which was the best in supporting MNV infection and replication, was used to generate HuhCD300lf VP2 by infection with this MNV VP2-expressing lentiviral vector. The lentiviral vector with the Huh7.5.1/CD300lf clone12-2 was further selected with 5 µg/mL puromycin for cells transduced with MNV VP2 and puromycin expression cassettes.

### Determination of infectivity

The infectivity of wild-type MNV produced from RAW264.7 or Huh7.5.1/CD300lf cells was determined by 50% tissue-culture infectious dose (TCID_50_) assay. RAW264.7 cells were seeded into 96-well plates (approximately 1 × 10^4^ cells/well) and inoculated with fivefold pre-diluted virus. After incubation at 37 ℃ with 5% CO_2_ for 4–5 days, infectivity was analyzed by virus-induced cytopathic effect. Virus titers were determined using the Spearman-Karber method.

### Production of recombinant MNV by DNA transfection

Plasmid constructs expressing MNV genes were transfected into 293T cells to generate recombinant viruses. Briefly, 250 ng of each pMNV_S7F_ derivative harboring mutations or deletions in ORF3 region and an equal amount of either pORF3 plasmid were mixed with 1 μL of P3000 solution (Thermo Fisher Scientific) in 50 μL of Opti-MEM (Thermo Fisher Scientific) and then combined with 50 μL of Opti-MEM supplemented with 1.5 μL of Lipofectamine 3000, and incubated for 20 min at room temperature. Before addition of Lipofectamine-DNA complex, approximately 42% of culture medium, 0.5 mL in 1.2 mL, of the 293T cells grown in 12-well plates was changed to the fresh medium containing 10 mM HEPES adjusted to pH 7.5. After transfection, cells were incubated for 48 h, and the cell-debris in the culture medium was removed by centrifugation at 12,000 ×*<u>g</u>* for 10 min at 4°C, and further cleared by filtration through a 0.2-micron filter, to be used as the source of the recombinant virus.

### Detection of the *infection* events by the mutant viruses

RAW264.7 or RAWVP2 cells infected with either wild-type or the mutant MNV were fixed at 48 hours post-inoculation (hpi) with 4% paraformaldehyde and were suspended in PBS with 0.1 % Triton-X. After blocking with 1% BSA/PBS, VPg and VP1 expression was detected by immunostaining using guinea pig anti-VPg antiserum and rabbit anti-VP1 antiserum, followed by Alexa488-conjugated anti-guinea pig IgG and Alexa594-conjugated anti-rabbit IgG secondary antibodies (Thermo Fisher Scientific), respectively.

Huh7.5.1/CD300lf and HuhCD300VP2 cells were fixed similarly at 48 hpi with 4% paraformaldehyde and suspended in PBS containing 0.1% Triton-X. Samples were blocked with 1% BSA/PBS, and the N-terminal protein expression was observed by immunostaining with anti N-terminal protein antiserum using epifluorescence microscope (BZ-X 800; KEYENCE). Image of entire well was taken by concatenating nine images of each well and positive cells were counted using the built-in image processing software of the BZ-X 800. The percentage of infected cells was calculated by dividing the number of cells expressing the N-terminal protein by the total number of DAPI-positive cells. Products from each construct at each passage were evaluated in independent five wells.

HuhCD300VP2 cells expressing fluorescent proteins (UnaG or Venus) upon infection with reporter viruses were detected using live-cell imaging with an epifluorescence microscope. Furthermore, Venus proteins were detected simultaneously with viral proteins after cell fixation. The DNA was stained with Hoechest33342. NanoLuc chemiluminescence, carried by the Nluc reporter virus recovered at 72 hours after infection, was detected according to the manufacturer’s instructions (Nano-Glo Luciferase Assay System, Promega) using a luminometer (EnSight Multimode Plate Reader, PerkinElmer).

Cell viability was determined using a CellTiter96 MTT assay kit (Promega). HuhCD300VP2 was incubated with the supernatant collected at each passage for 48 hours, and the extent of formazan product conversion from MTT by the cellular enzymes was analyzed by measuring the OD_490_ value.

### Purification of N-Nluc reporter particles and wild-type MNV

The pooled culture supernatant was centrifuged for 10 min at 10,000 ×*g*. The collected culture supernatant was treated with FBS-free DMEM containing 0.5% Zwittergent at room temperature for 10 min. Propagated viruses were pelleted through a 30% (wt/vol) sucrose cushion for 2 hours at 124,000 ×*g* using a Beckman SW32 Ti rotor, and further purified by isopycnic CsCl-gradient centrifugation in FBS-free DMEM (0.44 g/mL) for 24 hours at 150,000 ×*g*, using a Beckman SW55 Ti rotor. After centrifugation, the viruses, which were visualized as a white band near a density of 1.35 g/mL were collected using a micropipette. This solution was diluted 10 times in FBS-free DMEM, and the viruses were recovered by ultracentrifugation for 3 hours at 150,000 ×*g*.

### Confirmation of viral component proteins

Purified wild-type MNV in 2x Sample Buffer (0.125 M Tris-HCl, pH 6.8, 4% SDS, 10% sucrose, 0.01% bromophenol blue, 10% ß-mercaptoethanol) was used as the starting material for separation on 4–20 % Mini-PROTEAN TGX Stain-Free Protein Gels (Bio-Rad). The purified virus was loaded into the sample well and was run on the gel placed in a Mini-PROTEAN Tetra Vertical Electrophoresis Cell for 30 min at 200 V, 20 mA. The gel was imaged using the stain-free application on the ChemiDoc MP (Bio-rad) imager.

### Sample preparation for electron microscopy

Purified virions (10 µL) were adhered for 60 s onto freshly glow-discharged (to make the grid hydrophilic) carbon-coated copper grids. Excess sample was removed using the edge of a Whatman no. 4 filter paper wedge. While still wet, the grids were washed three or four times successively with a drop of 1% aqueous uranyl acetate each time, and the final drop was left on the grids for 40–45 s. Excess stain was removed using a filter paper wedge, and the grids were allowed to air dry and stored in a desiccated condition until further use for electron microscopy. The samples were viewed under a JEM2200FS (JEOL) cryoelectron microscope at an apparent magnification of 80,000 ×.

### Detection of viral RNA present in the cell-supernatant by Nested RT-PCR

The wild-type or mutant MNVs in the supernatant of 293T or RAWVP2 cells were immunoprecipitated by rabbit anti-MNV VLP antibody and Ab-Chapcher Mag (ProteNova). The precipitated MNV particles were eluted with 0.1 M glycine-HCl (pH 2.8) and immediately neutralized with 1 M Tris-HCl (pH 8.0). The eluate was purified for viral RNA using a QIAamp Viral RNA Mini (QIAGEN) and treated with Baseline-ZERO DNase (Epicenter) to digest potentially contaminating plasmid DNA. The viral RNA was reverse transcribed by SuperScript III First-Strand Synthesis System for RT-PCR (Thermo Fisher Scientific) with Tx30SXN primer, 5′-GACTAGTTCTAGATCGCGAGCGGCCGCCCTTTTTTTTTTTTTTTTTTTTTTTTTTTTTT-3′. For the 1^st^ amplification reaction Tx30SXN and MNV-F1, 5′-GCCATGCATGGTGAAAAG-3′ were used as the primers, and MNV-S, 5′-CCGCAGGAACGCTCAGCAG-3′, and MNV-R2, 5′-CAACCACCTTGCCAGCAG-3′, were used for nested PCR to detect MNV RNA. The PCR products were separated on 1.5% agarose gel, and the bands were visualized using ethidium bromide. The relative intensity of the bands was analyzed using ImageJ software.

To confirm that the deletions of the ORF3 regions were maintained during the passage of the mutants, virions in the supernatant were similarly immunoprecipitated and viral RNA was detected using the following set of primers for the nested PCR; MNV6401-6420 FW, 5′-TCGACTGTGCCCTTCCACAG-3′, and MNV7330-7308 RV, 5′-TCACAAAAGGTTTCTCTTCCAAC-3′.

### Antisera of MNV

Each polyclonal rabbit antiserum to MNV-S7 N-terminal protein (N-term), NTPase, VPg, RdRp, VP1 was produced by *Escherichia coli*-expressed and purified N-term, NTPase, VPg, RdRp, VP1 immunization to rabbit and guinea pig, respectively. Anti-MNV-S7 VP2 antiserum was produced with a truncated form of VP2 expressed in *E. coli* that removed the N- and C-terminal disorder regions. The truncated form of VP2 was purified and immunized into rabbit and guinea pig, respectively. Anti-MNV-S7 VLP antiserum was produced by recombinant baculovirus-expressed and purified MNV-S7 VLPs by CsCl sedimentation and immunized into rabbit and guinea pig.

## Conflicts of Interest

The authors declare no conflict of interest.

## Author Contributions

Conceptualization, K. K. and A. N.; Methodology, R.I., K.O., K.Y., R. T. T., A. K., Ku. K., K. K., and N. A.; Validation, R. I., K.O., and K. Y.; Formal Analysis, R. I., K.O., and K. Y..; Investigation, R. I., K.O., A. K., Ku. K. and K. Y.; Resources, K. K. and A. N.; Data Curation, R. I., K.O., and K. Y.; Writing – Original Draft Preparation, I. R., K. H., K. K., and A. N.; Writing – Review and Editing, I. R., R. T. T., K. H. K. K.; Visualization, I. R., and R. T. T.,.; Supervision, K. H., K. K., and A. N.; Project Administration, K. K. and A. N.; Funding Acquisition, K. K. and A. N.

## Acknowledgments and funding sources

Prof. Kazuyoshi Murata and Dr. Chihong Song, National Institutes of Natural Sciences, kindly provided the electron micrographs of recombinant virus.

## Funding

This study was supported by grants from the Japan Agency of Medical Research and Development under the Numbers 17fk0108034h, 18fk0108034h, 19fk0108034h, and 20fk0108121h to A. N. and K. K.

